# Dynamic [^18^F]Fluoroleucine PET Detects Impaired Cardiac Leucine Uptake Before Hypertensive Left Ventricular Hypertrophy Develops

**DOI:** 10.64898/2026.05.12.724048

**Authors:** William Terrell, Jie Li, Damodara N. Kommi, Megan Burt, Maurits A. Jansen, Shivashankar Khanapur, Susanna R. Keller, Bijoy Kundu

## Abstract

**Purpose:** Left ventricular hypertrophy (LVH) is a major complication of chronic hypertension and an independent risk factor for cardiovascular morbidity and mortality. There are currently no clinically validated markers available to identify hypertensive individuals at risk for developing LVH. In hearts of hypertensive rats, we previously described metabolic changes that precede LVH development, including in branched-chain amino acid (BCAA) metabolism. This study investigated whether cardiac leucine uptake, measured with dynamic 5-[^18^F]fluoroleucine positron emission tomography-computed tomography ([^18^F]FLE-PET/CT), was impaired and could serve as an *in vivo* marker for hypertension-induced LVH development.

**Procedures:** We synthesized [^18^F]FLE following established radiochemistry protocols and performed dynamic [^18^F]FLE-PET/CT imaging in 3-month-old spontaneously hypertensive rats (SHR) and normotensive Wistar-Kyoto (WKY) control rats (n = 4 per group). Cardiac magnetic resonance (CMR) imaging was conducted on the same animals for structural co-registration. A dual-output reversible two-tissue compartment model with spill-over (SP) and partial volume (PV) corrections was developed to quantify the first-pass rate constant (K_1_) and total distribution volume (V_t_ = K_1_/k_2_) for [^18^F]FLE. Protein expression of L-type amino acid transporter 1 (LAT1) and branched-chain keto acid dehydrogenase (BCKDH) phosphorylation status were assessed by immunoblotting of isolated heart tissue.

**Results:** SHR demonstrated markedly lower first-pass leucine uptake rates (K_1_) and total distribution volumes (V_t_) compared with WKY rats, consistent with reduced cardiac BCAA uptake. Concurrently, LAT1 (SLC7A5) expression was significantly reduced in SHR hearts compatible with decreased leucine uptake. Elevated BCKDH phosphorylation at Ser293 in SHR hearts indicated diminished BCKDH enzymatic activity and impaired BCAA catabolism.

**Conclusions:** Dynamic cardiac [^18^F]FLE-PET imaging successfully detects decreased leucine uptake in hypertensive rat hearts at 3 months of age, before LVH is established at 5 months. Reduced cardiac leucine uptake may thus serve as a surrogate marker for impaired cardiac BCAA metabolism and early in vivo indicator of cardiometabolic dysfunction that precedes LVH. The imaging approach holds translational potential for identifying hypertensive patients at risk for LVH progression.

## Introduction

Hypertension is a highly prevalent chronic condition affecting approximately 50% of the adult population in the United States^1, 2^. Among hypertensive individuals, approximately one-third develop left ventricular hypertrophy (LVH), a structural adaptation of the heart to chronic pressure overload that substantially increases the risk for major cardiovascular complications including heart failure, myocardial ischemia, atrial and ventricular arrhythmias, cerebrovascular disease, and dementia ^3–6^. The risk for complications correlates directly with the degree of LVH ^5, 6^, making early identification of individuals at-risk for developing LVH a critical clinical priority.

Despite the well-established complications of hypertension-induced LVH, no diagnostic markers are currently available to identify at risk hypertensive patients. LVH is typically not diagnosed until significant cardiac dysfunction is clinically apparent. It is then detectable by electrocardiography (ECG), echocardiography, or cardiac magnetic resonance imaging (CMR), with CMR considered the gold standard ^7^. Blood pressure control reduces but does not eliminate LVH and associated complications ^8^, underscoring the urgent need for biomarkers that allow detection of developing LVH at a stage when preventive or targeted therapeutic intervention are effective.

The heart has little storage capacity. With fluctuating energy demands cardiac substrate uptake is thus highly regulated ^9, 10^. The healthy heart relies primarily on fatty acids and glucose as substrates for energy production. Branched-chain amino acids (BCAAs) (leucine, isoleucine, and valine) contribute only approximately 1–2% to total cardiac ATP production under normal conditions ^11^. However, when elevated, BCAAs and their metabolic intermediates impair glucose and fatty acid metabolism and activate the growth-promoting signaling complex mTORC1 (mammalian target of rapamycin complex 1) that promotes cardiac hypertrophy ^12, 13^.

A growing body of evidence implicates cardiac metabolic dysfunction in the pathogenesis of pressure overload-induced cardiac hypertrophy. Previous work from our group using in vivo dynamic 2-[^18^F]-fluoro-2-deoxy-D-glucose (FDG) PET imaging and cardiac metabolomics in spontaneously hypertensive rats (SHR) demonstrated that chronic pressure overload leads to significant cardiac metabolic abnormalities in glucose, fatty acid, and BCAA metabolism as early as 2 months of age, when hypertension is established but overt LVH has not yet developed^14^. Metformin treatment of young hypertensive SHR normalized cardiac metabolic changes, improved cardiac function, and prevented LVH, supporting a causal relationship between metabolic dysfunction and LVH ^15^.

To summarize our previously published work ^14^, longitudinal dynamic FDG-PET imaging studies of SHR and WKY from 1 to 5 months of age, showed increased myocardial glucose uptake rates in SHR at 2 months together with elevated mean arterial pressure (MAP). CMR imaging showed reduced left ventricular ejection fraction (EF) beginning at 2 months in SHR, while statistically significant increases in the left ventricular mass/body weight (LVM/BW) ratio were observed only at 5 months (p < 0.05 SHR vs. WKY). At 3 months, LVM/BW was not yet significantly different between groups (p = 0.116). Metabolomics analysis of SHR hearts at 2 months revealed pronounced abnormalities in glucose, fatty acid, and BCAA metabolites. Related to BCAA metabolism, cardiac BCAAs and BCAA-derived carnitines were significantly elevated in SHR, and plasma BCAA levels were higher in SHR compared with WKY (p < 0.05). mTORC1 activity, as assessed by extent of phosphorylation of p70S6K on Thr389, was elevated in SHR hearts beginning at 2 months. Our findings suggest that in SHR, chronic pressure overload initiates cardiac metabolic abnormalities concurrently with the onset of hypertension but prior to detectable LVH.

To test whether changes in cardiac BCAA uptake contributed to observed changes in cardiac BCAA metabolites, we applied dynamic 5-[^18^F]fluoroleucine ([^18^F]FLE)-PET imaging of hearts. [^18^F]FLE is a fluorine-18 labeled leucine analog that is a promising tracer for measuring cardiac BCAA uptake non-invasively in vivo. It enters cells via the L-type amino acid transporter 1 (LAT1 or SLC7A5), a sodium- and pH-independent facilitative transporter ^16^ that primarily mediates BCAA uptake into cardiomyocytes ^17^. A prior study used [^18^F]FLE-PET in a small animal breast cancer model and evaluated the kinetics of the tracer *in vivo* ^17^, but did not compute kinetic parameters. [^18^F]FLE-PET has so far not been used to quantify cardiac BCAA uptake. We hypothesized that cardiac leucine uptake, as quantified by dynamic [^18^F]FLE-PET imaging, could serve as a proxy for impaired BCAA metabolism and an in vivo marker for cardiometabolic abnormalities that precede hypertension-induced LVH.

The present study describes: (1) the development and optimization of dynamic cardiac [^18^F]FLE-PET imaging and quantification in rats, using a dual-output reversible compartment model with spill-over (SP) and partial volume (PV) corrections; and (2) the application of this imaging approach to assess cardiac leucine uptake in SHR versus normotensive WKY controls, in conjunction with CMR imaging, and analysis of the expression of BCAA transporter LAT1 and phosphorylation of branched-chain keto acid dehydrogenase (BCKDH) as a readout for BCAA transport and catabolism. The ultimate goal is to establish cardiac leucine uptake as a translatable imaging biomarker for LVH risk stratification in hypertensive individuals.

## Materials and Methods

### Animals

All animal procedures were conducted in accordance with the National Institutes of Health (NIH) Guide for the Care and Use of Laboratory Animals and were approved by the Institutional Animal Care and Use Committee (IACUC) of the University of Virginia. Male spontaneously hypertensive rats (SHR) and normotensive Wistar-Kyoto (WKY) control rats (n = 4 per group) were studied at 3 months of age. SHR develop hypertension by approximately 2 months of age, with statistically significant LVH emerging at 5 months ^14^. All rats were maintained under standard housing conditions (12-hour light/dark cycle, ad libitum access to standard chow and water).

### Radiosynthesis of 5-[^18^F]Fluoroleucine ([^18^F]FLE)

The radiolabeling was performed following Chin et al ^17^ with significant modifications for a higher yield.

### Materials and Reagents

No-carrier-added aqueous [^18^F]fluoride ion was produced *via* the [^18^O(p,n)^18^F] nuclear reaction (Siemens Eclipse™ HP 11 MeV Cyclotron). The compounds mesylate precursor (precursor for [^18^F]FLE) and 5-fluoroleucine (reference standard for [^18^F]FLE) were custom synthesized from Jubilant Biosys Limited, India. Acetonitrile (99.8 %), Acetonitrile-d3 (≥ 99.8 %), anhydrous potassium carbonate (99.99 %), 4,7,13,16,21,24-hexaoxa-1,10-diazabicyclo[8.8.8]hexacosane (Kryptofix®222, 98 %), trifluroacetic acid (99 %), and HPLC-grade acetonitrile (≥ 99.9 %) were procured from Sigma-Aldrich, Inc. Furthermore, ethanol 200 proof USP was purchased from Decon Laboratories, Inc. and Gibco™ PBS, pH 7.4 solution was purchased from Thermo Fisher Scientific, Inc. and used as supplied.

All reactions were carried out in closed Thermo Scientific™ conical reacti-vial™ (5 ml capacity). Sep-Pak® light (46 mg) Accell™ plus QMA carbonate (part no. 186004540) and Sep-Pak™ plus short tC18 (part no. WAT036810) were purchased from Waters Technologies Corporation, Inc. Deionized water (18.2 MΩ·cm; ELGA PURELAB Flex 3) was used throughout the process. Radiochemical isolation and purification was conducted by semi-preparative radio-HPLC system (Logi-CHROM ONE from Lablogic Systems, Inc.) comprising of a low pressure gradient quaternary Pump, Manual injection valve (6-port/3-channel), a solvent mixer, a single wavelength UV-Detector (λ=254 nm) and Flow-Count radio-HPLC NaI detection system. Quality control analytical radio-HPLC was performed on a UFLC Shimadzu HPLC system equipped with a Low Pressure Gradient Binary Pump, a dual wavelength UV detector and a positron radiodetector (Posi-RAM 6; Lablogic Systems, Inc.). Radioactivity measurements were made with a CRC-55tW radioisotope dose calibrator (Mirion Technologies (Capintec), Inc.).

### Radiosynthesis Procedure

Aqueous [^18^F]fluoride (typically 12-14 GBq in ∼2.5 ml) was trapped on a Sep-Pak® light (46 mg) Accell™ plus QMA carbonate cartridge. The trapped [^18^F]fluoride anion was eluted into the reaction vial using 1 ml of a 95 : 5 (v / v) acetonitrile : water mixture containing K_2_CO_3_ (1.1 mg, 8.0 µmol) and Kryptofix 222 (5.8 mg, 15.4 µmol). The [K(K222)]^+^[^18^F]F^-^ complex was azeotropically dried under a stream of nitrogen gas (5 psi) at 110° C with the addition of 2 x 0.5 ml anhydrous acetonitrile. After cooling, a solution of the mesylate precursor (5 mg or 9.4 µmol) in anhydrous acetonitrile-d3 (0.6 ml) was added before the reaction vial was sealed and heated at 110° C for 15 minutes. After cooling for 2 min at room temperature, the crude reaction mixture was diluted with water (4.4 mL) and purified by gradient semi-preparative radio-HPLC using a Gemini C18 column (5 μm, 110 Å, 250 × 10 mm). The mobile phase consisted of 0.1% trifluoroacetic acid in water (TFA; solvent A) and acetonitrile (solvent B). The gradient was as follows: 0–1 min, 95% A/5% B; 1–12 min, linear gradient to 5% A/95% B; 12–15 min, 5% A/95% B; 15–20 min, 95% A/5% B. The flow rate was 5 mL/min, and UV detection was performed at 254 nm. The protected [^18^F]FLE fraction was isolated with a retention time of between 15.5-16.0 minutes and diluted with 25-30 mL water and passed through a Sep-Pak™ plus short tC18 cartridge to trap the protected [^18^F]FLE intermediate. The cartridge was washed with 5 mL of water, and the intermediate was eluted with 0.5 mL of TFA into a separate vial. The vial containing the TFA and the protected [^18^F]FLE intermediate was heated at 80°C under a stream of nitrogen until dry, effectively removing the Boc and tert-butyl ester protecting groups, and yielding the final product, [^18^F]FLE (**Figure 1**). The [^18^F]FLE was reconstituted in phosphate-buffered saline (PBS; 2.8 ml) and ethanol (0.2 ml) and subjected to quality control analysis before use in subsequent biological experiments. Radiosynthesis was performed at the University of Virginia (UVA) Radiochemistry Core and will in the future be automated on Elixys FLEX/CHEM + PURE/FORM and GE Tracerlab FX2 N synthesis modules to improve reproducibility, reduce radiation exposure, and enable GMP-compliant production for future translational studies.

**Figure 1.**
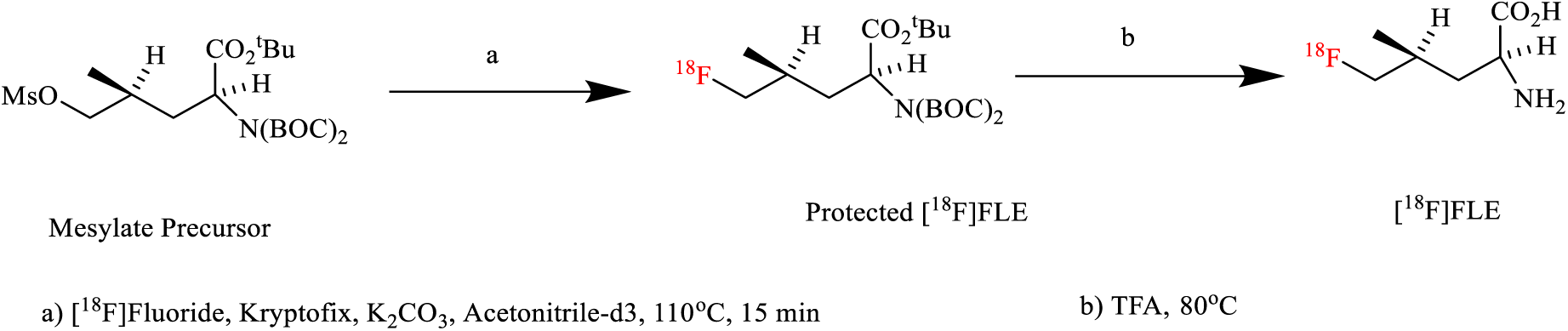
Schematic for synthesis of 5-[^18^F]-fluoroleucine

### Dynamic [^18^F]FLE-PET/CT Imaging

Rats were fasted for 5–6 hours prior to imaging. Under isoflurane anesthesia, a tail-vein catheter was placed and PET imaging (60-minute list-mode acquisition) using an Albira Si Trimodal PET/SPECT/CT scanner (Bruker Biospin, Ettlingen, Germany), a high-sensitivity, high-resolution small-animal PET imager was initiated coincident with slow intravenous injection (∼10 seconds) of 200–300 μCi [^18^F]FLE ^18^. Cardiac respiratory and heart rates were measured continuously using a small animal gating/monitoring system (Small Animal Instruments, Inc., model 1025L, Stony Brook, NY). Core body temperature was measured via a rectal probe and maintained at 37° Celsius with a regulated heating system. A 3-bed CT scan was obtained immediately following the PET scan for CT-based attenuation correction. List-mode data were sorted into 23 time frames (11 × 8 s; 1 × 12 s; 2 × 60 s; 1 × 180 s; 8 × 400 s) and sinograms reconstructed with attenuation, scatter, and random corrections using a Maximum Likelihood Expectation Maximization (MLEM) algorithm with 6 iterations and 0.5 mm isotropic voxel resolution.

### Cardiac Magnetic Resonance (CMR) Imaging

CMR was performed on the same rats on a separate day subsequent to PET to enable dynamic PET-MR co-registration and parametric PET quantification. All CMR studies were performed using a 9.4 T preclinical horizontal-bore BioSpec MRI system (Bruker Biospin, Ettlingen, Germany). Under anesthesia, ECG leads were placed subdermally for electrocardiogram (ECG) triggering. Bright-blood cine images were acquired in short-axis, vertical, and horizontal long-axis orientations using an ECG-triggered, respiratory-gated gradient echo sequence (TR/TE ≈ 7/1.6 ms; flip angle: 15°; field of view: 50 × 50 mm; matrix: 180 × 180; 2 signal averages; 24 time frames). Nine consecutive 1.5-mm-thick short-axis slices were acquired from base to apex of the left ventricle.

### PET-MR Image Co-Registration and Segmentation

PET-MR image co-registration was performed using PMOD image registration and fusion software (PFUS, version 3.9, PMOD Technologies Ltd., Zurich, Switzerland). CMR data were registered to CT using a manual linear registration with affine transformations (9 degrees of freedom: rotation, scaling, and translation) to generate a transformation matrix. This matrix was then applied to the dynamic PET data space. Myocardial volume and LV blood pool volumes of interest (VOIs) were segmented from high-resolution CMR images using 3D Slicer software. Segmentations were initially drawn on cine images and redrawn on T1 CMR images to ensure anatomical consistency. Image-derived input function (IDIF) was extracted from the LV blood pool VOI.

### Dual-Output Reversible Compartment Model for [^18^F]FLE Kinetics

A multi-parameter dual-output reversible two-tissue compartment model was developed and optimized to simultaneously estimate a model-corrected input function (MCIF) and myocardial kinetic parameters from dynamic [^18^F]FLE-PET images. This approach accounts for spill-over (SP) contamination and partial volume (PV) recovery effects inherent to small-animal cardiac PET imaging. The model was adapted from our previously validated dual-output non-reversible model for dynamic [^18^F]FDG-PET ^19^.

The dual-output model optimized the following composite objective function: O(p) = O_1_(p) + O_2_(p) + O_3_(p) (1) where O_1_ minimizes the sum of squared differences between model output and PET-derived time-activity curves (TACs) for both the blood (IDIF) and myocardial regions; O_2_ minimizes the squared error between model-predicted and PET-derived peak values; and O_3_ minimizes discrepancy between the MCIF and a late-time-point venous blood sample. The parametric blood input function Ca(t) was derived in parallel with myocardial tissue TACs. SP and PV corrections were incorporated through recovery coefficients and SP contamination factors (S_mb_, S_bm_, r_b_, r_m_) ^19^.

The reversible two-tissue compartment model was implemented as:

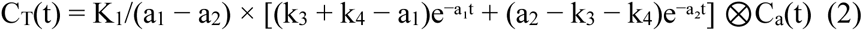

where K_1_–k_4_ are the kinetic rate constants, ⊗denotes convolution, and a_1_, a_2_ are derived from the kinetic parameters. Kinetic parameters computed included K_1_ (first-pass rate constant, in mL/min/g), V_t_ = K_1_/k_2_ (total distribution volume), and K_i_ = K_1_.k_3_/(k_2_ + k_3_) (net influx rate constant). Model fitting in Python by minimizing the objective function (Eq 1) was performed using the scipy.optimize module from the SciPy library on a 12-core Linux workstation using parallelized computation across time frames.

### Protein Expression Analysis

Rats were fasted for 6 hours and euthanized under 100% CO_2_ anesthesia with rapid dissection of hearts. Hearts were immediately frozen in liquid nitrogen, and tissue homogenates were prepared for immunoblotting. Briefly, aliquots of powdered frozen hearts were homogenized in Mammalian Protein Extraction Reagent (MPER; Thermo Scientific) containing Halt protease and phosphatase inhibitor cocktail (Thermo Scientific). Protein concentrations were determined using a bicinchoninic acid (BCA) assay (Pierce, Thermo Scientific). SDS samples (40 µg total protein per lane) were separated by SDS-PAGE (10% Tris-HCl polyacrylamide gels), transferred onto nitrocellulose membranes by electroblotting, and immunoblotted as described^25^. Specifically, membranes were blocked in 5% milk/Tris-buffered saline (TBS)/0.1% Tween-20 (TBST) for 2 h at 4°C, followed by overnight incubations at 4°C with antibody LAT1 (SLC7A5) (Invitrogen, Cat.# PA5-76141,1:1000), or antibody total BCKDH-E1 alpha (Cell Signaling Cat. #90198,1:1000) or antibody phospho-BCKDH-E1 alpha (Ser293) (Cell Signaling Cat. # 40368,1:1000) in 5% bovine serum albumin (BSA)/TBST. Immunoblotting conditions for phospho-p70S6K Thr389, a marker for mTORC1 activity, were as previously described ^15^. Membranes were subsequently incubated with HRP-conjugated secondary antibodies (GE Healthcare; Cat# NA934; 1:2,500) in 5% milk/TBST for 2 h at room temperature. Signals were detected using chemiluminescence and band densities determined with ImageJ software (NIH). The membranes were then stripped for 15 min using Restore Western blot stripping buffer (Thermo Scientific; cat. no. 21059), blocked in 5% milk TBST for 1h at 4°C, and incubated with beta actin antibodies (Santa Cruz, Cat.# sc-47778 HRP, 1:2,500) for 2 h at room temperature. Signals were detected using chemiluminescence and band densities determined with ImageJ software (NIH). All signal intensities were normalized to beta actin expression.

### Statistical Analysis

All data were analyzed using paired two-sample t-tests. p-values < 0.05 were considered statistically significant.

## Results

### Radiochemistry

[^18^F]FLE was prepared with a non-decay corrected radiochemical yield of 12 ± 2 % within 95-105 min (*n* = 5) from aqueous [^18^F]fluoride (**Figure 1**). The radiochemical purity of each delivered [^18^F]FLE batch was >99 %. The radiochemical identity of the [^18^F]FLE was confirmed by co-elution of the product with an authentic sample of [^19^F]FLE using analytical radio-HPLC using a Phenomenex Synergi™ LC column (4 µm, Hydro-RP 80 Å, 150 mm x 4.6 mm) (**Figure 2, top**). The mobile phase consisted of 0.1% trifluoroacetic acid in water (solvent A) and acetonitrile (solvent B). The gradient was as follows: 0 min (Initial), 95% A/5% B; 0–5 min, linear gradient to 5% A/95% B; 5–8 min, 5% A/95% B; 8–10 min, 95% A/5% B. The flow rate was 1 mL/min, and UV detection was performed at 220 nm (**Figure 2, bottom**). The retention time of authentic [^19^F]FLE and [^18^F]FLE was 3.3 minutes. The solution stability of formulated [^18^F]FLE was assessed over 3h at room temperature (*n* = 2) during which time no change in radiochemical purity was observed. To enable consistent high-throughput production for longitudinal studies, automation of the [^18^F]FLE radiosynthesis on an Elixys FLEX/CHEM+PURE/FORM and GE Tracerlab FX2 N modules will be implemented in the future.

**Figure 2.**
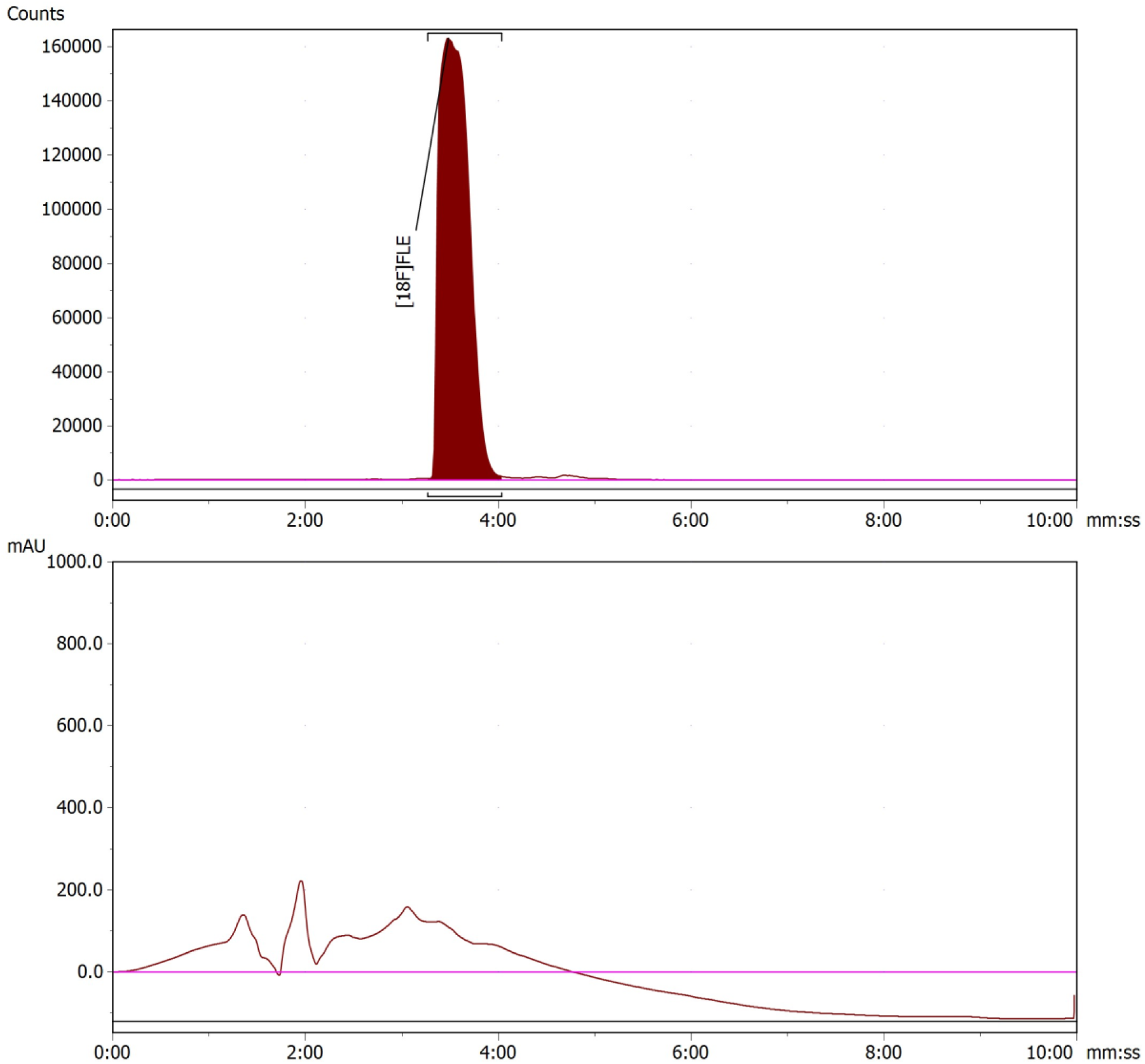
Radio (top) and UV (bottom) chromatograms of the final product 5-[^18^F]-fluoroleucine ([^18^F]FLE)

### Development and Validation of Dynamic Cardiac FLE-PET Imaging

Dynamic [^18^F]FLE-PET imaging of rat hearts was performed in 3-month-old male SHR (n=4) and WKY rats (n=3) using the Albira Si PET/SPECT/CT scanner. Time-activity curves (TACs) for both the LV blood pool (IDIF) and myocardial regions of interest were extracted from the reconstructed dynamic PET image series. The dual-output reversible compartment model was implemented, and a model-corrected input function (MCIF) was computed incorporating SP and PV corrections. Three objective functions (O_1_–O_3_) were simultaneously minimized to optimize model fit to both the IDIF TAC and myocardial TAC, with O_3_ incorporating a late-time-point venous blood sample as a tail-fitting constraint to suppress an artifactual rapid exponential drop in the MCIF. PET-MR co-registration using PMOD PFUS software was performed for all imaged animals. CMR-derived myocardial and LV blood pool segmentations provided accurate VOIs for subsequent parametric PET analysis. Parametric maps of K_1_, V_t_ = K_1_/k_2_, and K_i_ were generated for each animal (**Figure 3**). Computation of parametric maps was completed using parallelized processing across 12 CPU cores.

**Figure 3.**
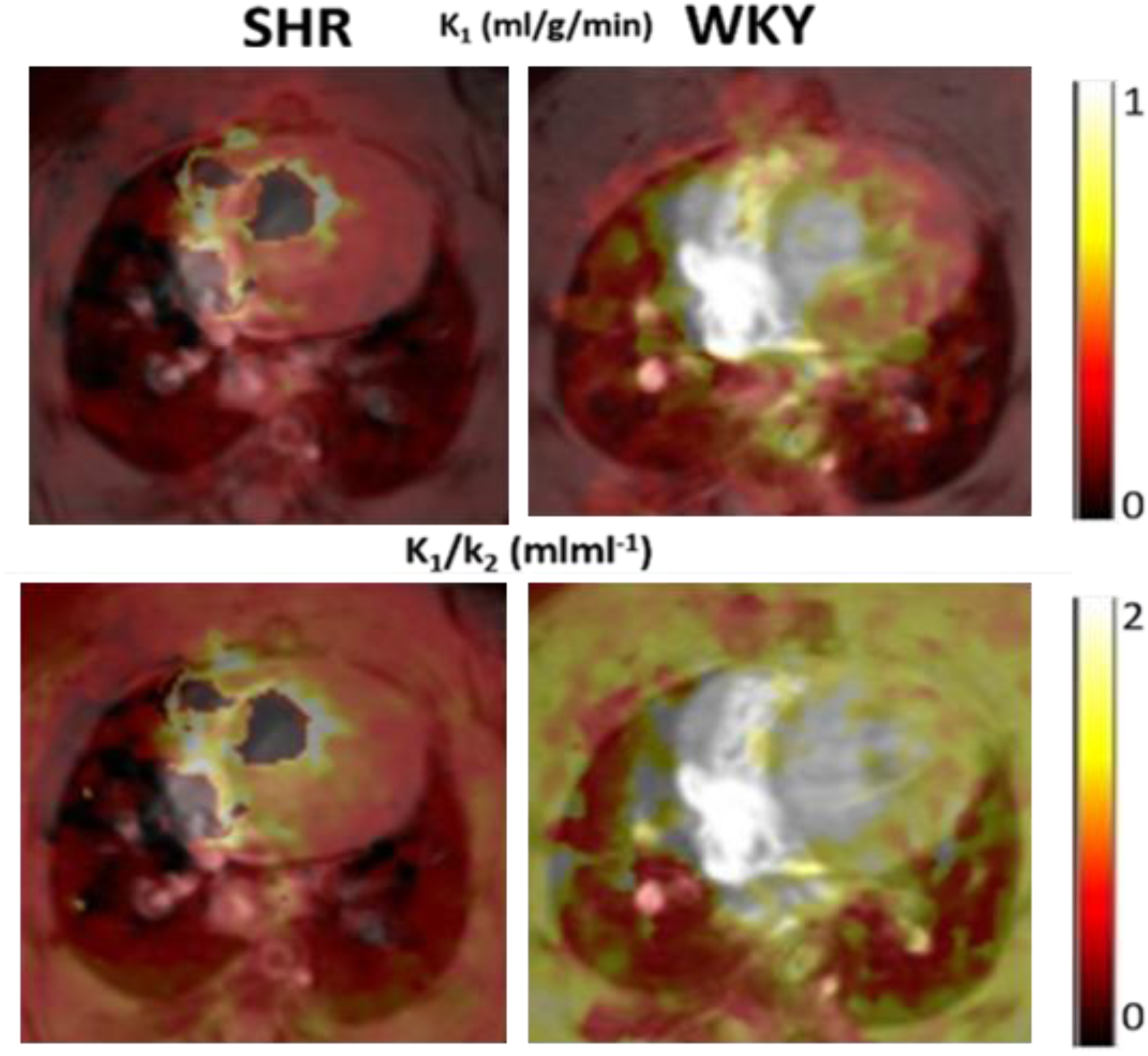
Parametric [^18^F]FLE PET (co-registered with CMR) demonstration of reduced [^18^F]FLE uptake rates in SHR (left panels) versus WKY hearts (right panels). SHR indicates spontaneously hypertensive rats; WKY, Wistar Kyoto rats.

### Decreased Cardiac Leucine Uptake in Hypertensive SHR

WKY rats demonstrated significantly higher first-pass leucine uptake rate constant (K_1_) and total distribution volume (V_t_ = K_1_/k_2_) compared with SHR at 3 months of age, consistent with decreased cardiac leucine uptake in SHR hearts. Despite the small sample size, the difference in K_1_ and V_t_ reached statistical significance (p = 0.019 and p = 0.042 respectively), demonstrating high sensitivity of [^18^F]FLE. The magnitude and direction of the effect were marked with ∼2.4 and 2.7-fold lower K_1_ and V_t_, respectively (**Figure 4**). No significant between-group differences were observed in k_3_ or net influx K_i_ in these preliminary studies, likely attributable to the rapid washout kinetics of [^18^F]FLE and absence of radiometabolite corrections in this initial analysis. Such limitations will be addressed in a fully powered future study through arterial blood sampling and radiometabolite correction.

**Figure 4.**
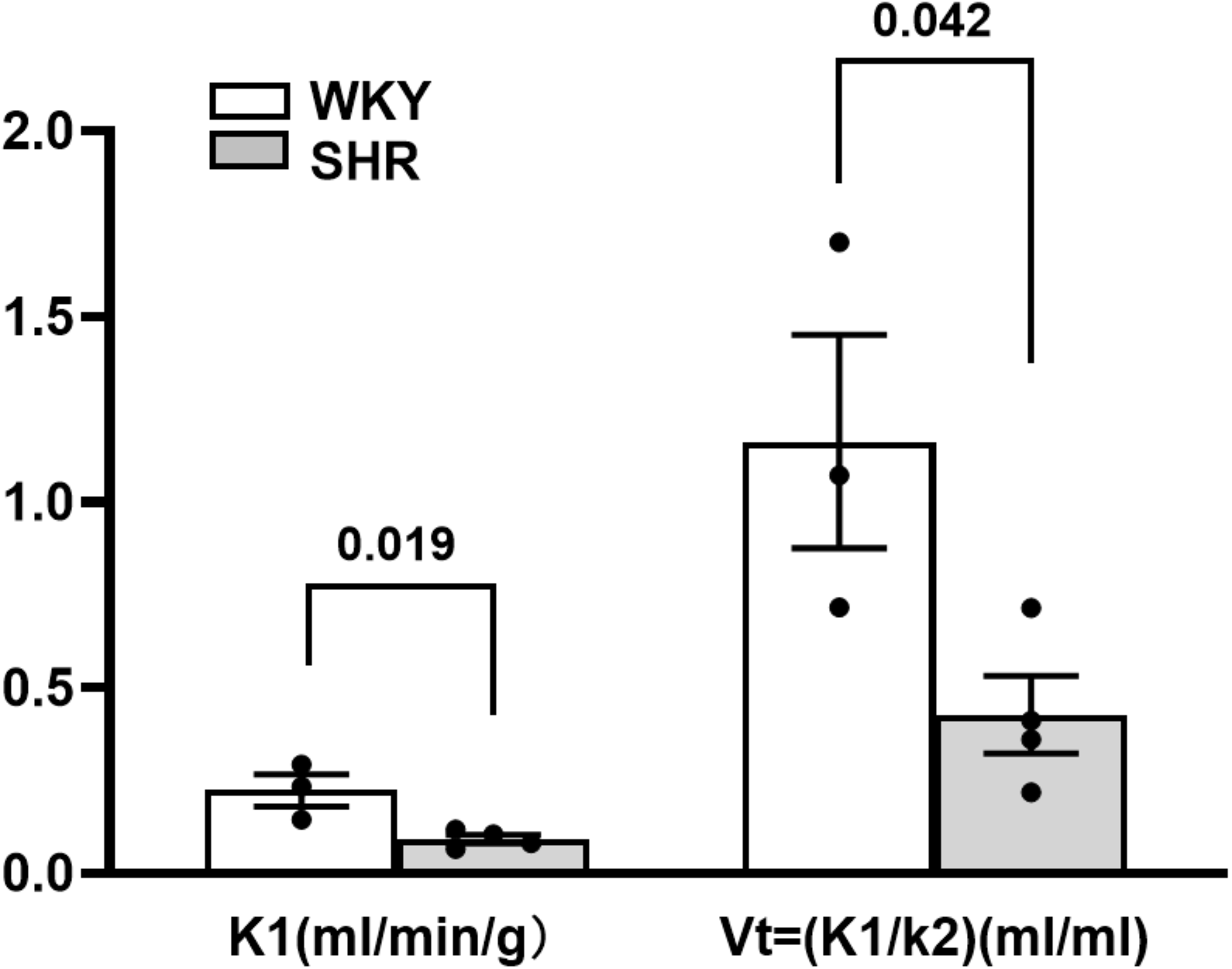
Computed rate constants for SHR and WKY with dynamic [^18^F]FLE PET. First pass rate constant K_1_ and ratio K_1_/k_2_ demonstrated reduced uptake in SHR (n=4) compared to WKY (n=3). Individual data points are shown for each strain with mean and standard error. SHR indicates spontaneously hypertensive rats; WKY, Wistar Kyoto rats. P<0.05 for both K_1_ and V_t_ (SHR vs WKY) indicating statistical significance.

### Changes of Key Proteins in BCAA Uptake and Metabolism in SHR Hearts

Above-described results demonstrated impaired cardiac leucine uptake. To test whether the major BCAA transporter LAT1 (SLC7A5) was changed, we immunoblotted heart homogenates with SLC7A5 antibodies. **Figure 5A** shows that in hearts of the 3-month-old SHR used for [^18^F]FLE-PET imaging, SLC7A5 is significantly reduced (by ∼30%), consistent with decreased [^18^F]FLE uptake. Immunoblotting analysis of previously harvested hearts ^14^ demonstrated that SLC7A5 expression was also significantly decreased (by ∼20%) in SHR hearts at 2 months of age when SHR are hypertensive, but no cardiac structural changes are evident. SLC7A5 expression remains lower in SHR hearts (by ∼30%) at 5 month of age when LVH is apparent. At 1 month of age, when SHR are pre-hypertensive, SLC7A5 expression in SHR hearts was similar to control WKY (**Figure 6**), suggesting that SLC7A5 downregulation only occurs when SHR are hypertensive.

**Figure 5.**
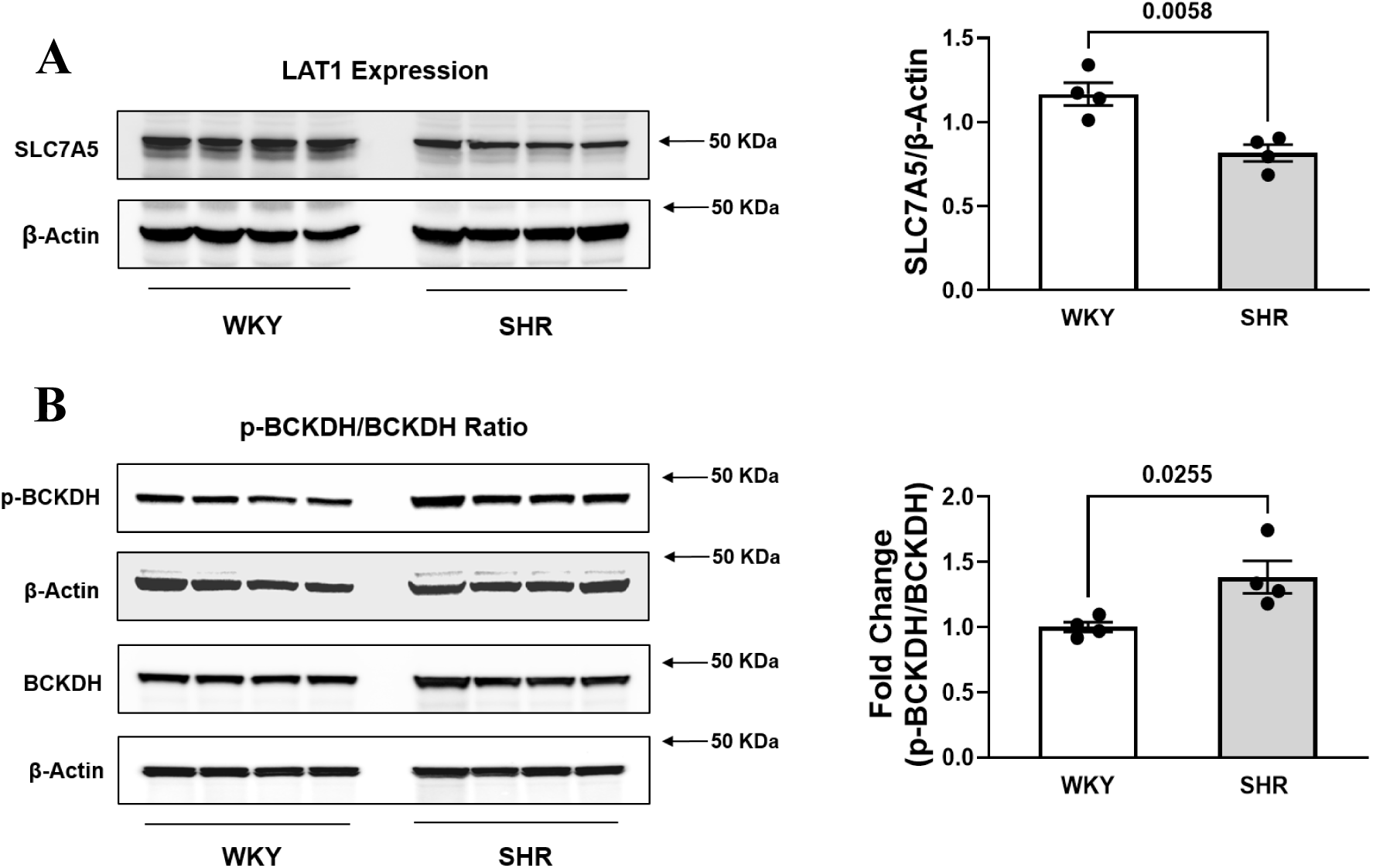
LAT1 (SCL7A5) and p-BCKDH in 3-month-old SHR (n=4) and WKY (n=4) hearts collected after [^18^F]FLE PET Imaging. **A.** LAT1 (SLC7A5) and β-actin immunoblots (left) and quantification of LAT1 (SLC7A5) expression normalized to β-actin (right). **B.** Immunoblots for p-BCKDH (p-Ser293), total BCKDH, and β-actin (left), and quantification of the ratio of p-BCDKH to total BCKDH each normalized for β-actin (right). Individual data points are shown for each strain with mean and standard error. SHR indicates spontaneously hypertensive rats; WKY, Wistar Kyoto rats. P<0.05 (SHR vs WKY) indicates statistical significance.

**Figure 6.**
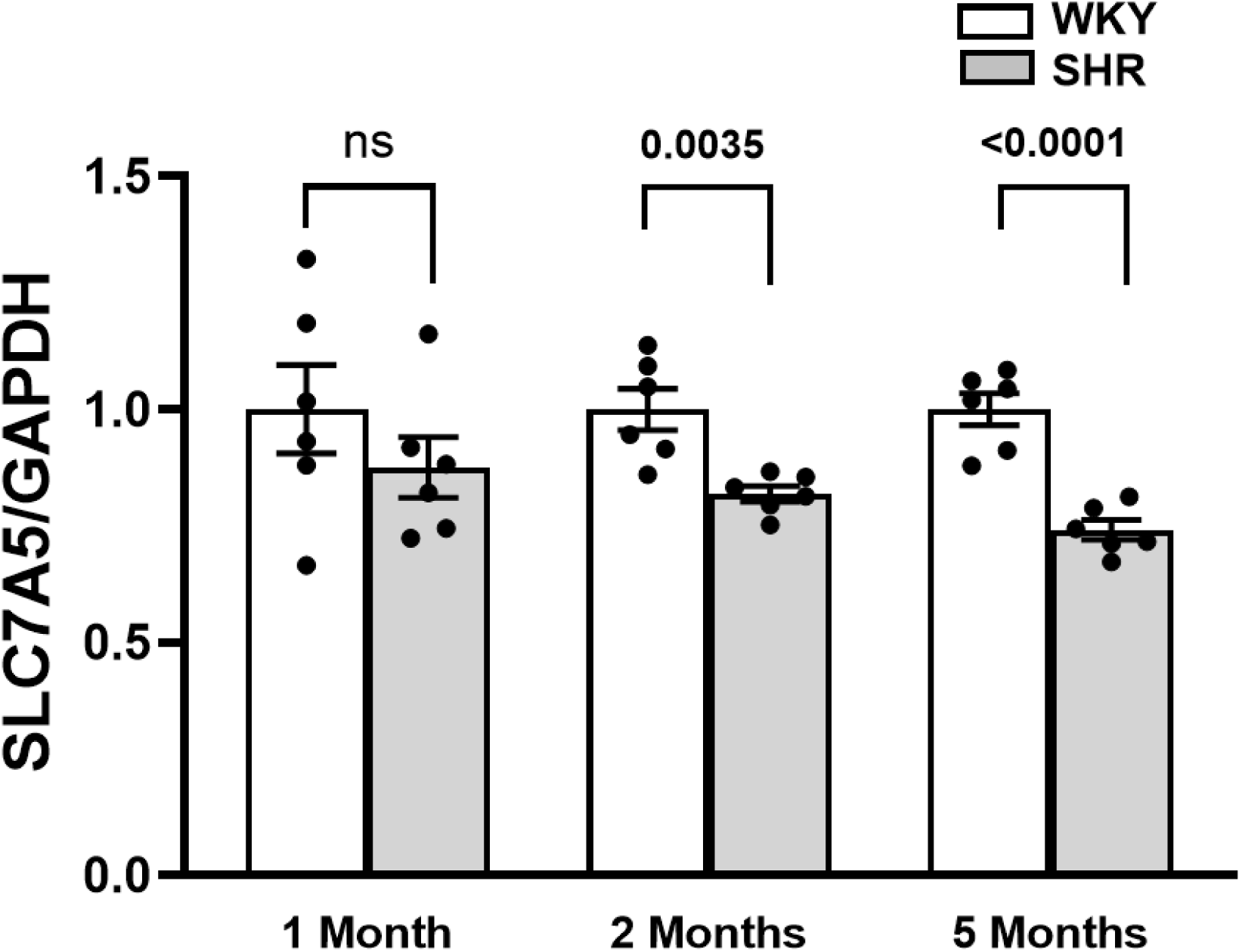
LAT 1 (SCL7A5) Expression in SHR. LAT 1 expression (normalized to GAPDH) in SHR (n=6) compared to WKY (n=6) at 1, 2, and 5 months of age from previously harvested rat hearts [12]. Individual data points are shown for each strain at each time point with mean and standard error. SHR indicates spontaneously hypertensive rats; WKY, Wistar Kyoto rats. P<0.05 at 2 and 5 months (SHR vs WKY) indicates statistical significance. ns: non-significant.

Above results showing decreased leucine uptake with decreased leucine transporter expression while, as we previously described in 2-month-old SHR, cardiac BCAAs and BCAA-derived carnitines are elevated^14^, suggested additional changes in intracellular BCAA metabolism. Following uptake, BCAAs are catabolized by branched-chain amino acid transferase (BCAT) to branched-chain keto acids (BCKAs) that are subsequently irreversibly decarboxylated by branched-chain keto acid dehydrogenase (BCKDH) to branched chain acyl-CoA derivatives. These derivatives are further processed to acetyl-CoA and succinyl-CoA that enter the Krebs cycle^20^. BCKDH activity is regulated by reversible phosphorylation on serine residue 293 (Ser293) by BCKDH kinase (BCKDHK), with increased phosphorylation reducing BCKDH activity. As shown in **Figure 5B**, BCKDH phosphorylation at Ser293 was markedly elevated in SHR hearts at 3 months of age with no significant change in total BCKDH protein abundance. Elevated BCKDH phosphorylation is consistent with decreased BCKDH activity leading to impaired BCAA catabolism and increased intracellular BCAAs and BCAA metabolites despite decreased BCAA uptake.

## Discussion

This study presents the first application of dynamic 5-[^18^F]fluoroleucine ([^18^F]FLE) PET imaging to quantify cardiac leucine uptake rates *in vivo* in a hypertensive rat model. Our key findings are: (1) [^18^F]FLE can be reliably synthesized and used for dynamic cardiac PET imaging in small animals; (2) a dual-output reversible compartment model successfully quantifies myocardial [^18^F]FLE kinetics with SP and PV corrections; (3) cardiac leucine uptake, as measured by K1 and Vt, is diminished in 3-month-old SHR prior to detectable LVH; and (4) decreased leucine uptake is accompanied by reduced BCAA transporter LAT1 (SLC7A5) expression and elevated BCKDH phosphorylation suggesting impaired intracellular BCAA catabolism.

We are currently unable to identify hypertensive individuals at heightened risk for developing LVH. LVH increases the risk for major cardiovascular events by 2–4-fold ^4^, and complications, including heart failure and arrhythmias, are associated with substantial morbidity and mortality ^6^. Current clinical tools detect LVH only once structural remodeling is advanced. A robust in vivo imaging marker that predicts LVH development would open paradigm shifting opportunities for the design of novel approaches to prevent or ameliorate complications of LVH.

Our laboratory has previously established that cardiac metabolic abnormalities, including altered BCAA metabolism, precede LVH in SHR ^14^. The present data extend these findings using [^18^F]FLE-PET to non-invasively measure leucine uptake, a surrogate for BCAA transport. The observed decrease in K1 and Vt in SHR hearts is consistent with downregulated LAT1 (SLC7A5) expression. LAT1, the predominant L-type amino acid transporter in heart ^16^, has higher leucine affinity compared to LAT2, making [^18^F]FLE uptake specifically reflective of LAT1-mediated transport capacity. Downregulation of LAT1 is consistent with a negative feedback loop whereby increased intracellular BCAA and BCAA metabolites downregulate LAT1 expression ^21^. A decrease in cardiac BCAA uptake may thus serve as a marker for impaired BCAA catabolism as well as increased cardiac growth triggered by increased BCAA and BCAA metabolites.

The choice of [^18^F]FLE as a PET tracer for cardiac BCAA metabolism is well-grounded. FLE is structurally and functionally homologous to leucine, the most abundant BCAA ^22^ and primary allosteric activator of mTORC1 signaling. Other amino acid analogs such as (2S,4R)-4-[^18^F]fluoroglutamine and 3-[^18^F]-L-α-methyl-tyrosine, respectively, have been successfully used in oncologic imaging for SLC1A5 and LAT1 (SLC7A5)-mediated transport ^23, 24^.

The dual-output reversible compartment model developed here represents a methodological advance for small-animal cardiac PET imaging. Standard semi-quantitative standardized uptake values (SUV) or simplified graphical methods (Patlak, Logan plots) lack the sensitivity and specificity required to capture the nuanced kinetic changes in tracer transport and distribution that characterize early-stage metabolic dysfunction. Our 15-parameter model simultaneously recovers the MCIF, corrects for SP and PV contamination, and extracts mechanistically informative kinetic parameters (K_1_, V_t_, K_i_) from a single dynamic acquisition. Incorporating a late-time-point blood sample as a tail-fitting constraint (O_3_ cost function) resolved the problem of artifactual MCIF decline observed in initial iterations of the model. This will be critical for enabling longitudinal studies without repeated arterial catheterization.

This study also has limitations. The preliminary imaging dataset was underpowered for formal statistical comparison, although we observed group differences in K_1_ and V_t_ reaching statistical significance. A fully powered study (n = 10 per group) is planned to address this. Radiometabolite corrections for the IDIF were not applied in the initial analyses, which may have introduced underestimation of true k_3_ and K_i_ due to blood input contamination. Since radiometabolite contamination typically contributes to later time points in IDIF compute, the results reported for first pass K_1_ and V_t_ may not be affected. To optimize model accuracy, ground-truth arterial blood sampling with radio-HPLC radiometabolite profiling will be incorporated in a future larger study. Additionally, the current dataset is limited to male SHR at a single time point. To provide a more comprehensive characterization of BCAA uptake dynamics relative to LVH development, future studies will include both sexes, multiple time points, and additional hypertensive rat models.

For early diagnosis to be relevant for prevention of LVH complications, therapeutic approaches to restore BCAA metabolism will need to be designed. A BCKDH kinase inhibitor (BT2) has recently been shown to improve BCAA metabolism in pressure-overload heart failure models ^25^. To demonstrate that cardiac [^18^F]FLE-PET imaging results reflect BCAA catabolic capacity, future experiments will test whether leucine uptake is normalized following BT2 treatment.

Cardiac [^18^F]FLE-PET imaging has potential for clinical application. Our group has previously demonstrated that dual-output compartment models developed for rodent cardiac [^18^F]FDG-PET can be adapted to dynamic human cardiac [^18^F]FDG-PET ^26^. Given declining costs of PET imaging, advances in total-body time-of-flight PET scanners ^27^, and the availability of deep-learning parametric reconstruction algorithms that reduce required radioactivity dose ^28, 29^, [^18^F]FLE-PET could be translated to humans with acceptable radiation burden. A practical clinical workflow might involve initial screening of hypertensive patients for plasma BCAA metabolite biomarkers, followed by [^18^F]FLE-PET imaging of a limited number of humans with and without the relevant metabolite signature to test for associated cardiac metabolic changes. A metabolite signature may subsequently suffice as a screening tool to identify patients at high risk for LVH or reduce the number of individuals requiring an additional [^18^F]FLE PET. This tiered approach would balance diagnostic precision with feasibility and cost-effectiveness.

## Conclusions

Dynamic [^18^F]FLE PET imaging successfully detects reduced cardiac leucine uptake in SHR prior to detectable LVH. Decreased leucine uptake, quantified as K_1_ and V_t_ through a dual-output reversible compartment model with SP and PV corrections, is accompanied by reduced LAT1 expression and elevated BCKDH phosphorylation, indicating impaired BCAA transport and catabolism in SHR hearts. The cardiometabolic perturbations precede LVH.

Our findings support the hypothesis that decreased cardiac leucine uptake could serve as an in vivo marker for impaired BCAA catabolism that triggers increased cardiac growth and cardiometabolic dysfunction preceding hypertension-induced LVH. With further development, cardiac [^18^F]FLE-PET imaging has the potential to become a diagnostic tool for identifying hypertensive individuals at greatest risk for progression to LVH who may benefit from early metabolically targeted interventions.

## Acknowledgements

This study was supported in part by NIH grant R01 HL 123627 and Radiology and Medical Imaging Start-up Funds (to BK). The study utilized [^18^F] FLE radiopharmaceutical synthesis and delivery services provided by the UVA Radiochemistry Core Facility supported by the UVA School of Medicine, RRID: SCR_025471 and UVA Molecular Imaging Core facility support with National Institutes of Health S10OD021672 funding for the Albira Si trimodal scanner and S10OD025024 funding for the Bruker 9.4T BioSpec MRI scanner. We thank Jeremy Gatesman at the UVA Center for Comparative Medicine for help with tail-vein catheterizations for tracer administration.

## Declarations

### Ethics Approval

All animal procedures were conducted in accordance with the NIH Guide for the Care and Use of Laboratory Animals and approved by the IACUC of the University of Virginia.

### Data Availability

Raw PET imaging data, compartment model code, and other datasets supporting the conclusions of this article will be made available upon reasonable request to the corresponding author.

